# Single-cell genome-wide association reveals a nonsynonymous variant in *ERAP1* confers increased susceptibility to influenza virus

**DOI:** 10.1101/2022.01.30.478319

**Authors:** Benjamin H. Schott, Liuyang Wang, Xinyu Zhu, Alfred T. Harding, Emily R. Ko, Jeffrey S. Bourgeois, Erica J. Washington, Thomas W. Burke, Jack Anderson, Emma Bergstrom, Zoe Gardener, Suzanna Paterson, Richard G. Brennan, Christopher Chiu, Micah T. McClain, Christopher W. Woods, Simon G. Gregory, Nicholas S. Heaton, Dennis C. Ko

## Abstract

Diversity in the human genome is one factor that confers resistance and susceptibility to infectious diseases. This is observed most dramatically during pandemics, where individuals exhibit large differences in risk and clinical outcomes against a pathogen infecting large portions of the world’s populations. Here, we developed scHi-HOST (single cell High-throughput Human in vitrO Susceptibility Testing), a method for rapidly identifying genetic variants that confer resistance and susceptibility to pathogens. scHi-HOST leverages scRNA-seq (single-cell RNA-sequencing) to simultaneously assign genetic identity to individual cells in mixed infections of cell lines of European, African, and Asian origin, reveal associated genetic variants for viral entry and replication, and identify expression quantitative trait loci (eQTLs). Applying scHi-HOST to influenza A virus (IAV), we identified eQTLs at baseline and in genes that are induced by IAV infection. Integration of scHi-HOST with a human IAV challenge study (Prometheus) revealed that a missense variant in *ERAP1* (*Endoplasmic reticulum aminopeptidase 1;* rs27895) was associated with IAV burden in cells and human volunteers. Functional studies using RNA interference, ERAP1 inhibitor, and overexpression of alternative alleles demonstrated that ERAP1 is exploited by IAV to promote infection. Specifically, the nonsynonymous substitution, which results in a glycine to aspartate substitution at ERAP1 residue 348, would disrupt the substrate binding pocket of ERAP1, likely resulting in a significantly altered preference for substrates, poorer catalytic efficiency, or both. Finally, rs27895 exhibits substantial population differentiation, with the higher frequency of the minor T allele in two African populations likely contributing to the greater permissivity of cells from these populations to IAV infection. scHi-HOST is an important resource for understanding susceptibility to influenza and is a broadly applicable method for decoding human genetics of infectious disease.

## Introduction

Infectious diseases have been a leading cause of mortality throughout human history. These past exposures to pathogens have conferred strong selective pressures on the human genome, driving adaptation as resistance alleles become more common. When a pathogen remains endemic to a region, resistance alleles continue to provide protection against the pathogen that drove selection, as demonstrated by malaria protection by the sickle cell allele (rs334) (Allison, 1954). Alternatively, episodic selection by epidemics and pandemics may leave lasting consequences on genomes for future infections, as well as other unexpected aspects of human health impacted by the same cellular pathways (Pittman et al., 2016). Identification of human genetic differences that regulate resistance and susceptibility to infectious diseases could reveal individuals and populations at risk for more severe infections and genes that could be targeted for novel therapies.

In pursuit of genetic differences that impact resistance and susceptibility, approaches using human populations have focused on identification of either common or rare variants. For influenza A virus (IAV), small, underpowered GWAS using a common variant approach have failed to identify any genome-wide significant loci (Chen et al., 2015; Garcia-Etxebarria et al., 2015; Zhou et al., 2012). Coupling small candidate association studies with functional evidence has revealed associations of common variants in *IFITM3* with severe influenza (Allen et al., 2017; Everitt et al., 2012). On the other end of the frequency spectrum, ascertainment of individuals with very severe disease and next-generation sequencing has identified rare variants predisposing to severe influenza in *IRF7* (Ciancanelli et al., 2015) and *TLR3* (Lim et al., 2019). However, challenges are associated with deconvoluting the mechanisms underlying identified common or rare variants. Complementary *in vitro* approaches for identifying human genetic variation can control for differences in exposure, pathogen genetic diversity, comorbidities, and access to medical care and can reveal mechanisms underlying genetic association. Such approaches could be used to broadly probe human genetic diversity for resistance and susceptibility to emerging pathogens and would be a powerful tool, especially as outbreaks develop in single geographic locations.

Previously, we developed a cellular GWAS platform, Hi-HOST (Hi-throughput Human in *vitrO* Susceptibility Testing), in which lymphoblastoid cell lines (LCLs; Epstein-Barr virus immortalized B cells) from hundreds of genotyped individuals are exposed to identical doses of a pathogen and assessed for cellular host-pathogen traits including entry (Alvarez et al., 2017; Bourgeois et al., 2021), replication (Wang et al., 2018), cell death (Ko et al., 2012; Ko et al., 2009; Salinas et al., 2014), and cytokine response (Barnes et al., 2022; Schott et al., 2020; Wang et al., 2018). This approach revealed human genetic differences associated with cellular traits that are also associated with infectious disease risk (Alvarez et al., 2017; Gilchrist et al., 2018; Malaria Genomic Epidemiology, 2019) and outcomes (Ko et al., 2012; Wang et al., 2017). Thus, while Hi-HOST screening with LCLs captures phenotypes measured from a single cell type, effects of SNPs that can be assayed in LCLs have been validated in other cellular models, animal models, and humans. Others have also carried out studies of cytokine levels (Li et al., 2016; Mikacenic et al., 2013) and of response eQTLs (reQTLs) following stimulation with various pathogens or pathogen-related molecular patterns (Barreiro et al., 2012; Fairfax et al., 2014; Lee et al., 2014; Nedelec et al., 2016). However, Hi-HOST and similar studies are labor- and time-intensive and can be skewed due to batch effects in measuring hundreds of individual cellular infections or exposures over a period of months or even years. Additionally, incorporation of new technology for simultaneous identification of eQTLs combined with GWAS of pathogen measurements could help fill the mechanistic gap from genetic variant to cellular infection phenotype.

Here, we present a rapid, high-throughput approach called scHi-HOST (single-cell Hi-HOST) that uses scRNA-seq for simultaneous identification of alleles associated with both gene expression changes and genetic resistance to IAV. This method leverages the genetic diversity of dozens of individuals from multiple populations in a single infection and revealed both phenotypic differences in IAV susceptibility among populations and a common missense variant in *ERAP1* (rs27895) associated with IAV burden in LCLs and a human flu challenge study. Coupling scHi-HOST with human challenge studies is a broadly generalizable approach for understanding human genetic susceptibility to infectious disease using a small number of individuals in a rapid timeframe.

## Results

### A single-cell approach for discovering genetic resistance to infection

We developed scHi-HOST to integrate the identification of common human genetic variants that impact gene expression and cellular infection traits into a single infection assay. scHi-HOST uses scRNA-seq for simultaneous genotypic assignment of thousands of individual cells and dense phenotyping of host and virus. The scHi-HOST pipeline has 4 major steps: 1) infection and scRNA-seq data generation, 2) assignment of individual cells to genotyped individuals, 3) GWAS of pathogen-based burden phenotypes, and 4) eQTL identification of host gene expression using allele-specific expression and total reads across genotypes (**Fig. 1A**). In this study, equal number of LCLs from 48 individuals across 3 populations (LWK (Luhya in Webuye, Kenya), GBR (British in England and Scotland), and CHS (Southern Han Chinese) from the 1000 Genomes Project (Genomes Project et al., 2015)) were mixed and infected for 24 hours with a laboratory adapted human H1N1 IAV strain (A/Puerto Rico/8/34, PR8). This experiment is referred to as scHH-LGC, based on the populations included. Consistent with previous reports of B cells being direct targets for IAV entry (Dougan et al., 2013; Lersritwimanmaen et al., 2015; Tumpey et al., 2000), LCLs were highly infectable by IAV. A scRNA-seq library was prepared using 10X Genomics Chromium Next GEM Single-Cell 3’ technology (Zheng et al., 2016). Mixed LCLs were frozen into aliquots that can be thawed and screened for future pathogens or other stimuli in <1 week.

**Figure 1.**
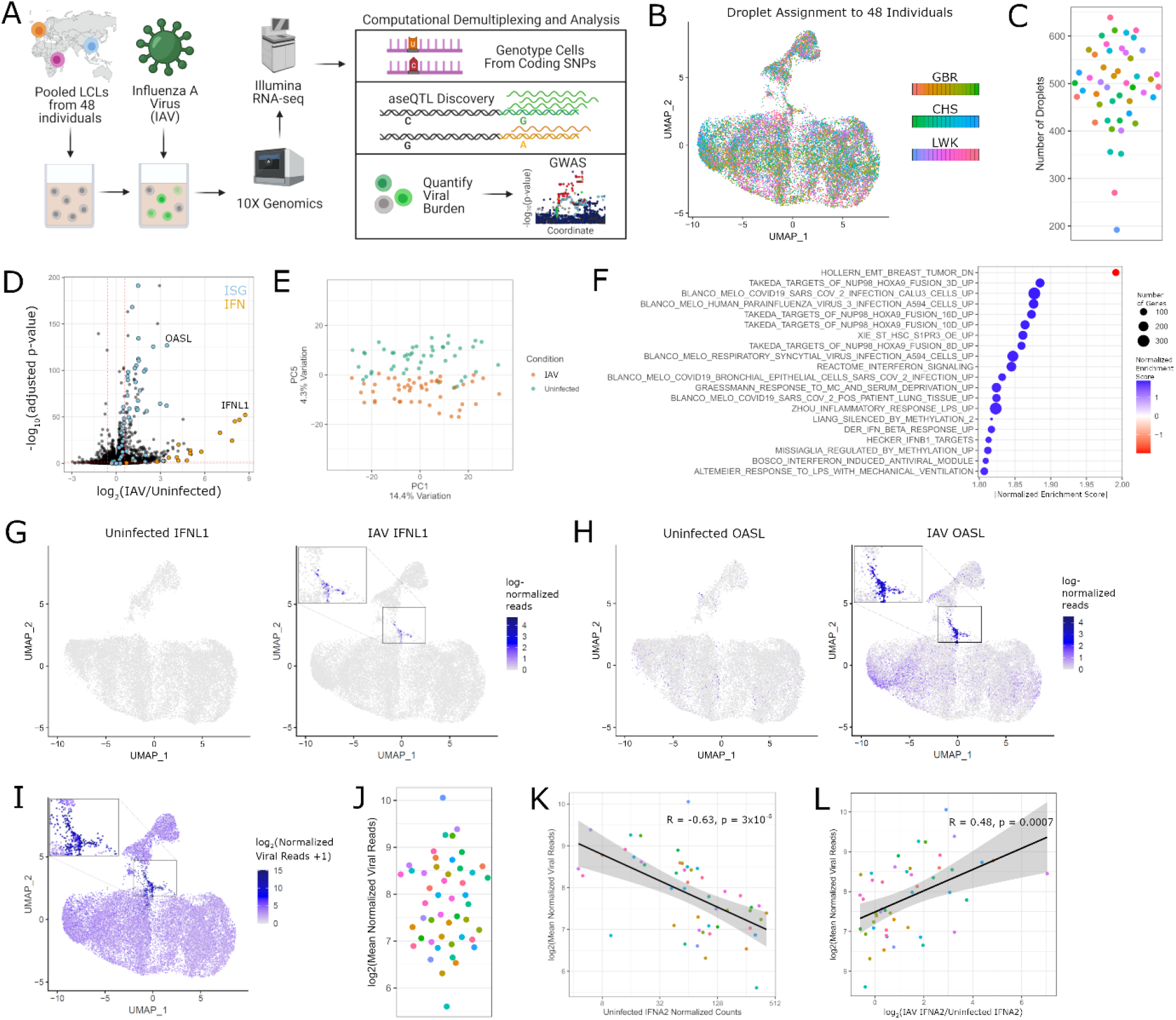
scHi-HOST is a rapid platform for cellular host-pathogen genome-wide association. **(A)** Flowchart of scHi-HOST with influenza A virus. **(B)** UMAP plot demonstrates assignment of individual cells to one of 48 genotyped LCLs for uninfected and IAV-infected scHi-HOST samples. Color-coded by shades for each population. (**C**) The number of individual cells measured for each LCL in the scRNA-seq analysis. Color-coding is the same as panel B. **(D)** Volcano plot of pseudo-bulk analysis of uninfected vs. infected LCLs revealed upregulation of IFNs and ISGs. Pseudo-bulk analysis was performed by aggregating all cells of the same LCL identity (total of 48 pseudo-bulk samples) and performing differential expression in DESeq2 (Love et al., 2014). Genes are color-coded by functional class; orange: Browne Interferon Responsive Genes (GSEA C2 geneset), blue: all 21 human interferon genes, black: all other genes. Dotted red horizontal line indicates p = 0.05. Dotted red vertical lines indicate fold-change > 1.5 or < 0.5 relative to uninfected. **(E)** Pseudo-bulk analysis shows uninfected transcriptomes and infected transcriptomes segregate along PC5. **(F)** GSEA shows upregulation of ISGs and other viral response gene sets. Plotted gene sets are top 20 absolute normalized enrichment score, all FWER p < 0.05. (**G**) UMAP plots of *IFNL1* in uninfected and IAV-infected LCLs. (**H**) UMAP plots of *OASL* in uninfected and IAV-infected LCLs. (**I**) UMAP plot demonstrates highly variable IAV burden across individual cells and a cluster of highly infected cells. UMAP plots were generated by combining uninfected and IAV samples for normalization and plotting each condition separately with Seurat (Butler et al., 2018). In contrast, uninfected cells had a total of 19.6 normalized viral reads, with reads detected in only 23 out of 20,129 cells. These likely represent index-hopped artifacts and not true contaminants. (**J**) Mean viral reads across 48 LCLs. Phenotype was generated by normalizing by total read depth per cell (human and viral reads) and aggregating all cells of the same LCL identity. (**K**) Negative correlation of baseline *IFNA2* and viral burden. *IFNA2* was chosen as it has the highest baseline expression, though similar results were observed with other expressed IFNα genes (Fig. S2A). Correlation coefficient and p-value from Spearman’s correlation. (**L**) Positive correlation of induced *IFNA2* and viral burden. Positive correlation between induction of gene expression and viral burden was similarly observed for other IFNα genes (Fig. S2B). Correlation coefficient and p-value from Spearman’s correlation.

Following next-generation sequencing, SNPs in the oligo-dT primed cDNA reads facilitated the unequivocal assignment of nearly all individual cells to one of the 48 genotyped LCLs using Demuxlet (Kang et al., 2018) (**Fig. 1B**; median number of individual cells for each LCL = 501, **Fig. 1C**). We achieved a singlet rate of 73% with 99.9% of singlets assigned to a single genotype with > 99% confidence.

LCLs displayed a transcriptional response similar to other IAV infected cell types (Li et al., 2019; Zhuravlev et al., 2020). Differentially expressed genes (DEGs) were dominated by upregulation of interferon (IFN) and interferon-stimulated genes (ISGs) (**Fig. 1D; Table S1**). PCA analysis showed convincing separation of uninfected and IAV infected LCLs along PC5, accounting for 4.3% of the variation (**Fig. 1E**). Gene set enrichment analysis (GSEA) (Subramanian et al., 2005) confirmed upregulation of ISGs along with multiple other gene sets for viral infection including multiple respiratory virus response gene sets, interferon-induced antiviral modules, and the inflammatory response (**Fig. 1F**, all FWER p < 0.05; **Table S2**).

At the single-cell level, we found that induction of IFN is widely variable and only detected in a minority of cells (**Fig. 1G**). In contrast, ISGs, including *OASL*, while variably expressed, are more broadly induced throughout the infected culture (**Fig. 1H**). Thus, while the antiviral response is robust, it is driven by a small number of cells that highly induce IFN to then induce ISGs more broadly.

To determine if this transcriptional heterogeneity can be explained by cell-to-cell differences in viral burden, we mapped and quantified viral transcripts, given that poly-adenylated IAV RNA was captured with the LCL transcriptome. A UMAP plot revealed that most cells have low but detectable levels of viral reads, but there is a cluster of high-burden cells with a distinct host transcriptional response (**Fig. 1I**). This high level of heterogeneity is consistent with previous work using scRNA-seq in IAV-infected A549 lung epithelial cells (Russell et al., 2018). We observed the most highly burdened cells in the culture expressed the highest levels of IFNs and ISGs in response to virus. Aggregating the IAV reads for each of the 48 LCLs revealed large differences, with the mean influenza reads varying ∼20-fold across different LCLs (**Fig. 1J**). It was noted that viral burden did not correlate with copy number of EBV (**Fig. S1A**), used by the 1000 Genomes Project in immortalizing the LCLs, or with the number of each LCL recovered in the scHi-HOST experiment (**Fig. S1B**). Thus, there is wide variation in viral burden across cells at the single-cell level and on aggregate across LCLs and these differences correlate with differences in the expression of important antiviral genes.

Comparison of gene expression in uninfected and infected LCLs vs. viral burden revealed that high IFNα expression at baseline in the uninfected state is correlated with low viral burden in infected cells at 24 hours (**Fig. 1K; Fig. S2A**). In contrast, after infection, high induction of IFNα is correlated with high viral burden (**Fig. 1L, Fig. S2B**). Thus, high baseline IFNα levels prior to infection appear to be protective; however, during infection, high levels of IAV replication, as evaluated by IAV genome copy number, lead to the highest levels of IFNα induction in individual cells. This is consistent with a previous study using single-cell RNA sequencing of IAV infected cells to link basal expression to susceptibility to infection (O’Neill et al., 2021).

In summary, in a single infection, we deconvoluted cells from 48 different individuals and observed highly variable flu burden and transcriptional phenotypes. These methods revealed baseline IFN levels contribute to resistance against influenza infection. We next turned to identifying human genetic differences that regulate cellular response to infection.

### Common human genetic differences impact transcriptional response to IAV in cells

We identified human SNPs associated with variation in gene expression in uninfected and infected cells using RASQUAL (Robust Allele-Specific Quantitation and Quality Control) (Kumasaka et al., 2016). RASQUAL tests for the association of SNPs vs. gene expression level by measuring both linear regression among genotypes and allele-specific expression in heterozygous individuals while accounting for sex biases and confounding factors. Observed - log(p-values) of the data deviated strongly from theoretical and empirical null distributions, indicating many true positives under both conditions (**Fig. 2A**). We compared the results of RASQUAL analysis on scHH-LGC with a second scHi-HOST experiment of 48 additional LCLs from ESN (Esan from Nigeria), IBS (Iberian from Spain), and KHV (Kinh from Vietnam) (referred to as scHH-EIK) and observed strong correlation between the two datasets (**Fig. 2B**). These datasets were combined to provide the greatest power in detecting eQTLs. The most significant eQTLs were detected in both uninfected and IAV-infected conditions (**Fig. 2C**).

**Figure 2.**
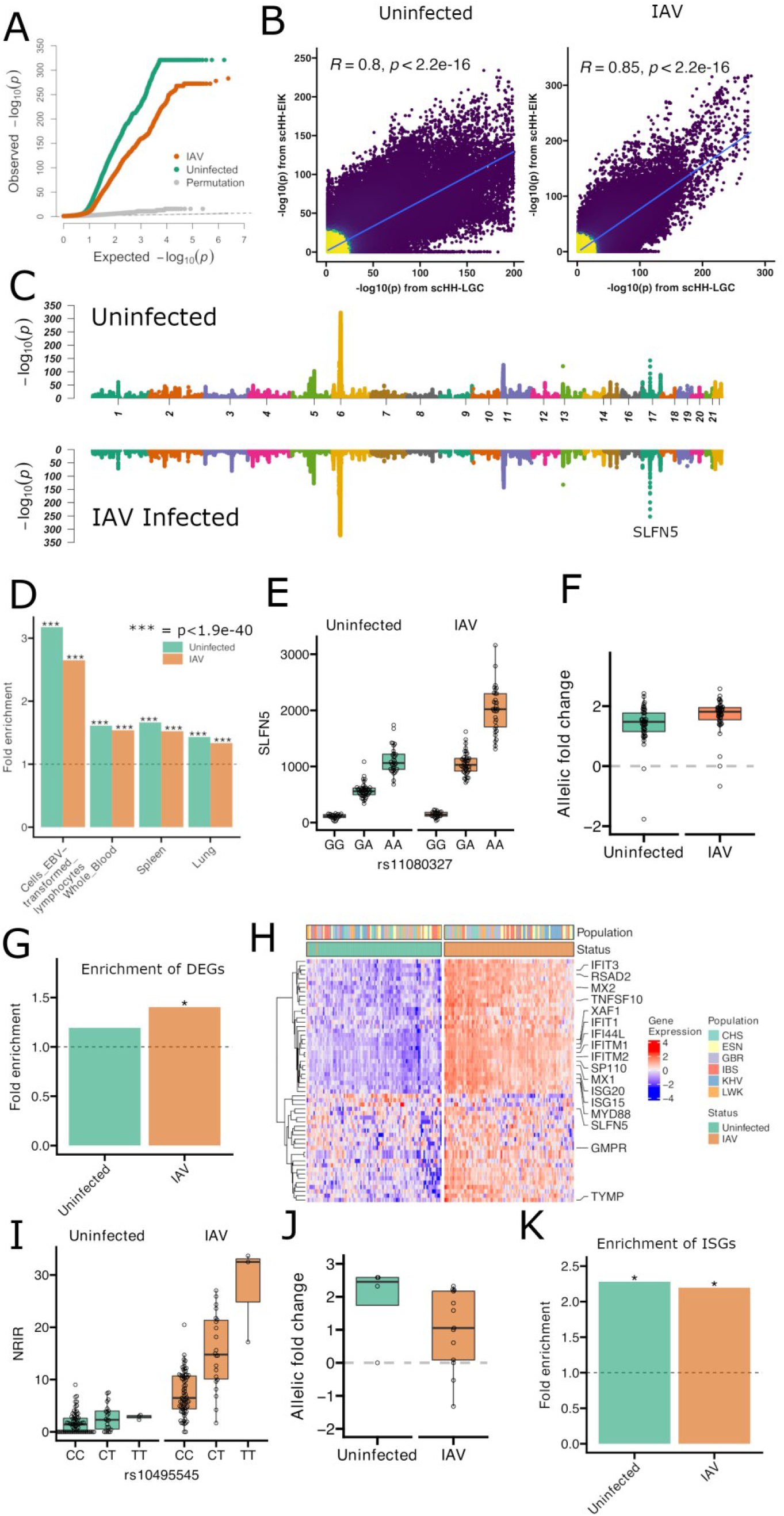
Common human genetic differences regulate transcriptional response during cellular influenza infection. **(A)** QQ plot of observed, permutated, and expected -log(p-values) for eQTLs demonstrates deviation of observed p-values from neutral expectation for both uninfected and IAV-infected LCLs. **(B)** Correlation of eQTLs identified by scHH-LGC and scHH-EIK. -log p-values for all SNP-Gene pairs with p < 0.05 in scHH-LGC were plotted against -log p for the same SNP-gene pairs in scHH-EIK. **(C)** Miami plot of cis-eQTLs for uninfected and IAV infected cells identified in the combined scHH-LGC + scHH-EIK dataset in 1MB windows using FDR-corrected p-values. **(D)** scHi-HOST eGenes from uninfected and IAV-infected LCLs were highly enriched in GTEx. scHi-HOST eGenes were defined using a 2-step FDR < 0.05 and Fisher’s exact test was used to calculate the significance of enrichment of scHi-HOST eGenes against GTEx eGenes (FDR < 0.01). (**E, F**) rs11080327, the most strongly associated cis-eQTL outside of the MHC region (Uninfected 2-step FDR = 5.3 x 10^-140^, IAV 2-step FDR = 1.3 x 10^-249^), is associated with expression of *SLFN5* based on linear regression of genotypic medians (E) and allelic imbalance for uninfected and infected LCLs (F). As *SLFN5* is an ISG, the expression of the gene is higher in IAV-infected LCLs and the association in that context is stronger. **(G)** IAV-induced DEG (differentially expressed genes; p < 0.05, fold-change > 1.5) are significantly enriched in IAV-infected eGenes. DEGs between uninfected and IAV-infected were identified using a pseudo-bulk approach as described in Method. Fisher’s exact tests were used to test the significance of overlap between DEGs and eGenes and calculate p-values. Strong enrichment is observed only with IAV infection (*, p = 6.8×10^-3^). **(H)** Heatmap of expression of the 56 DEGs (p < 0.05; absolute FC > 1.5) that are also eGenes. Numerous ISGs are labeled. LCLs are color-coded by population. (**I, J**) Genotypic median plots (I) and allelic imbalance plots (J) for the association of rs10495545 with *NRIR* (2-step FDR = 0.0003 for IAV-infected, 2-step FDR = 1.0 for uninfected). *NRIR* is induced 4.95-fold with IAV infection and is an IAV-specific eQTL. (**K)** Enrichment analysis of ISGs in eGenes from uninfected and IAV-infected LCLs. Strong enrichment is observed under both conditions, consistent with LCLs having baseline IFN production in the uninfected state (*, p < 0.001).

A 2-step approach was used to identify eGenes (unique genes associated with eQTLs) utilizing Benjamini-Hochberg correction for both the SNPs tested in each window and for the number of genes tested. For uninfected and IAV-infected LCLs at a 2-step false-discovery rate FDR < 0.05, we detected 2265 and 3326 eGenes, respectively (**Table S3**). Many of the eQTL associations replicated in the GTEx dataset (Consortium, 2020). We observed 3.17-fold enrichment (p=1.26×10^-117^) for uninfected eGenes in GTEx LCLs (GTEx FDR < 0.01) and 2.65-fold enrichment (p = 6.02×10^-140^) for IAV eGenes, representing 33.01% of the eGenes from the infected sample. Further, enrichment was observed across multiple tissues in GTEx, including whole blood (1.61-fold and 1.54-fold enrichment for IAV and uninfected eGenes respectively, both p<1.9×10^-40^), spleen (1.66-fold and 1.52-fold enrichment for IAV and uninfected respectively, both p<1.9×10^-40^), and lung (1.43-fold and 1.33-fold enrichment for IAV and uninfected respectively, both p<1.9×10^-40^) (**Fig. 2D**). Thus, as others have noted (Flutre et al., 2013), LCLs serve as a valid system for identification of eQTLs that are relevant in many tissues, and we have determined that eQTLs identified in LCLs with scHi-HOST are reproducible across other LCL datasets, related primary tissues, and even other tissues.

The most significant association outside of the HLA region in both uninfected and infected LCLs was the association of rs11080327 with *SLFN5* (**Fig. 2E, F**, Uninfected 2-step FDR = 5.3 x 10^-140^, IAV 2-step FDR = 1.3 x 10^-249^). Notably, the expression of this gene increased with IAV infection (1.80-fold; adjusted p = 0.004) (consistent with *SLFN5* being an ISG (Katsoulidis et al., 2010)) and the effect size of the SNP (allelic-fold change (Mohammadi et al., 2017)) was greater in the infected LCLs (**Fig. 2F**) (Uninfected aFC = 2.6; IAV-infected aFC = 3.6; p = 0.0001).

We observed an enrichment of IAV-induced genes (see Fig. 1D) in the IAV eGenes (1.4-fold enrichment; p = 6.8 x 10^-3^) but not in the uninfected eGenes (1.19-fold enrichment; p = 0.18), indicating that context-specific eQTLs can be uncovered by the scHi-HOST approach (**Fig. 2G**). In total, we detected 56 DEGs that were also eGenes, many of which were also ISGs (**Fig. 2H**). For example, we discovered an eQTL (rs10495545) for the long-ncRNA *NRIR* (*Negative Regulator of Interferon Response* (Kambara et al., 2014); also known as AC017076.5 and lncRNA-CMPK2). This lncRNA is induced 4.95-fold during IAV infection in LCLs and is an eQTL only in the IAV infected state (**Fig. 2I**; **2J**; p = 0.0003 for IAV-infected, p = 1.0 for uninfected). ISGs are enriched in both uninfected and IAV infected scHi-HOST datasets (**Fig. 2K**), consistent with LCLs having baseline expression of ISGs that is induced further during IAV infection.

### Integration of scHi-HOST and functional annotation reveals a nonsynonymous SNP in *ERAP1* that increases IAV susceptibility

While scHi-HOST is a powerful method for eQTL identification in response to pathogens, it also serves as a method for rapid association testing of functional subsets of SNPs. Given the modest size of our combined scHi-HOST dataset (n = 96), even associations that surpass genome-wide significance (traditionally p < 5 x 10^-8^) must be evaluated carefully for corroborating evidence. For the phenotype of mean IAV burden, a single SNP (rs1923314; p = 1.8 x 10^-8^) surpassed genome-wide significance (**Fig. S3**; full summary statistics available at Duke Research Data Repository https://doi.org/10.7924/r4g163n0s.). However, this SNP is in a gene desert and is not a cis-eQTL, making functional follow-up challenging. Therefore, we examined statistical evidence of association for SNPs that have functional consequences: cis-eQTLs as identified above and non-synonymous SNPs based on annotation.

Examination of stratified QQ plots (Schork et al., 2013), restricted to eQTLs (FDR < 0.05 in uninfected or IAV-infected lead eGene-eQTL; MAF > 0.1), demonstrated deviation from neutral expectation (**Fig. 3A**). rs12103519, associated with expression of *TNFSF12* (*Tumor Necrosis Factor Superfamily member 12*; Uninfected RASQUAL 2-step FDR = 0.001, IAV RASQUAL 2-step FDR = 0.02) had the strongest association with mean IAV reads (p = 1.3 x 10^-5^). Similarly, stratification of nonsynonymous variants also demonstrated deviation from neutral expectation (**Fig. 3B**), with a missense SNP (rs27895; G346D) in *ERAP1* (*endoplasmic reticulum aminopeptidase 1*) having the lowest p-value (p = 2.9 x 10^-5^; **Fig. 3C; Fig. S4**). We tested both *TNFSF12* and *ERAP1* for a role in regulating IAV infection by reducing expression with siRNA and assessing infection using a IAV mNeon reporter strain (Harding et al., 2017). While knockdown of *TNFSF12* had no effect (**Fig. 3D**), *ERAP1* siRNA in LCLs led to a consistent decrease in IAV infection, suggesting a proviral role (**Fig. 3E**; p = 0.01 in NA19020, p = 0.01 in NA19399). *ERAP1* encodes an aminopeptidase, with the canonical function of trimming N-terminal residues from peptides for MHC class I presentation (Saric et al., 2002; Weimershaus et al., 2013; York et al., 2002). The derived T allele of rs27895 is associated with higher mean IAV reads and encodes a G346D amino acid change. Notably both *ERAP1* and the related *ERAP2* have been reported to undergo IAV mediated changes in expression, but the functional consequences of *ERAP1* induction (Wang et al., 2013) and *ERAP2* alternative isoform usage (Ye et al., 2018) on IAV infection are unknown. To confirm that *ERAP1* is proviral, we treated the A549 lung epithelial cell line with an ERAP1 inhibitor (4-methoxy-3-(N-(2-(piperidin-1-yl)-5-(trifluoromethyl)phenyl)sulfamoyl)benzoic acid; (Maben et al., 2020)) and observed a dose-dependent reduction in the percentage of IAV-infected cells (**Fig. 3F**; p < 0.0001 at 10 µM).

**Figure 3.**
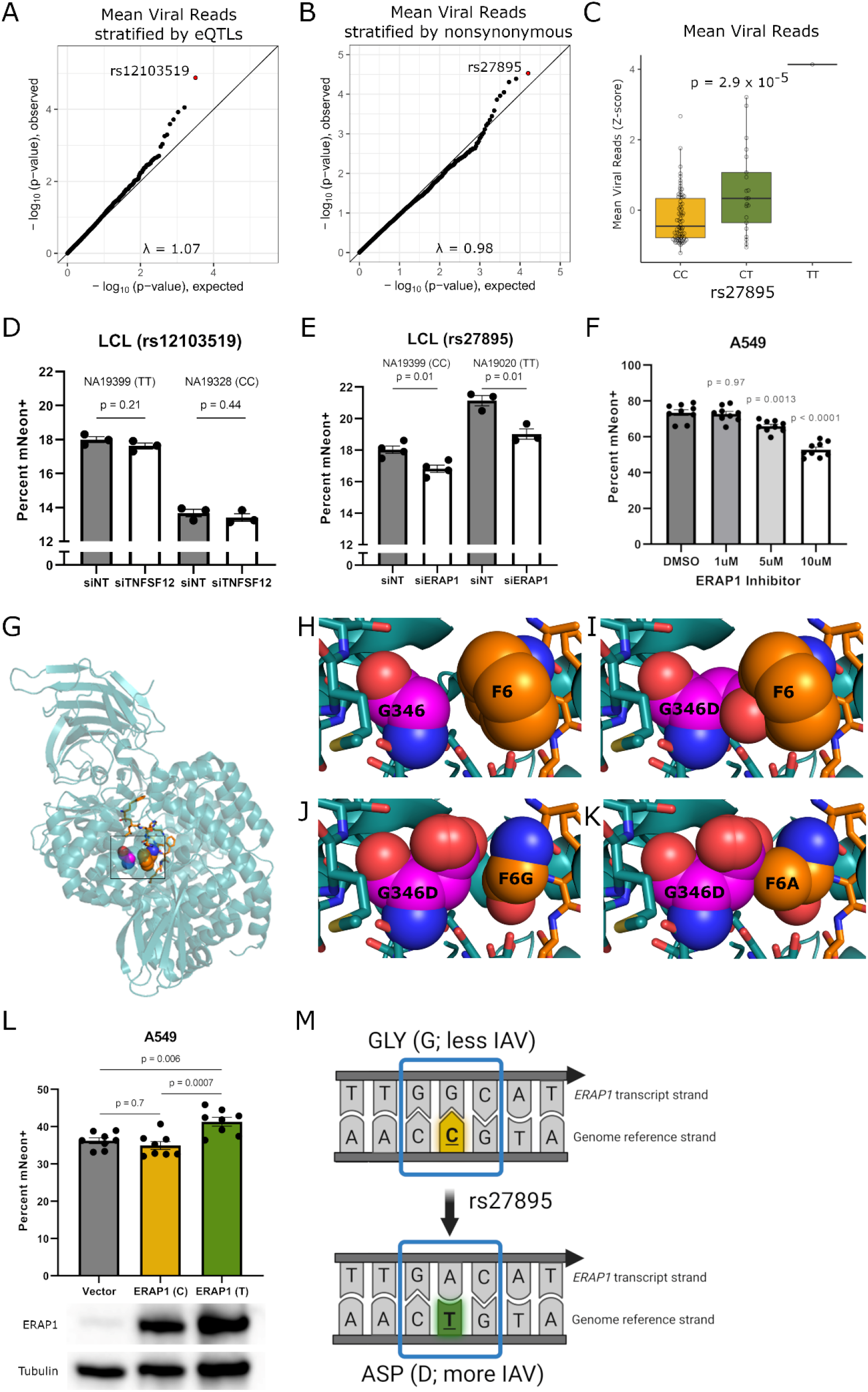
A nonsynonymous variant in *ERAP1* regulates IAV burden in cells. **(A)** Stratified QQ plot for association with mean IAV reads restricted to RASQUAL 2-step FDR eQTLs (FDR < 0.05; MAF > 0.1). Plotting eQTLs reveals a SNP associated with expression of *TNFSF12* that has a lower p value than expected by chance. **(B)** Stratified QQ plot for association with mean IAV reads restricted to nonsynonymous SNPs. Plotting nonsynonymous variants (MAF > 0.1) reveals a SNP in *ERAP1* that has a p value lower than expected by chance. **(C)** Genotype median plot for mean IAV reads as a function of rs27895 genotype. (**D**) RNAi against *TNFSF12* in either NA19399 (TT for rs12103519) or NA19328 (CC for rs12103519) demonstrates reducing *TNFSF12* expression (mean 87% knockdown +/- 4% (SEM) for NA19328 and mean 73% knockdown +/- 6% (SEM) for NA19399) does not significantly affect the percentage of IAV infected cells at 24 hours in either LCL. Replicates from 3 separate experiments were normalized to correct for between-experiment variation. P values are from unpaired t-tests. **(E)** RNAi against *ERAP1* in either NA19020 (TT for rs27895) or NA19399 (CC for rs27895) demonstrate that reducing *ERAP1* expression (mean 80% knockdown +/- 14% (SEM) for NA19020 and mean 70% knockdown +/- 8% (SEM) for NA19399) decreases the percentage of IAV infected cells at 24 hr in both LCLs. Replicates from 4 separate experiments were normalized to correct for between-experiment variation. One experiment involving NA19020 was excluded because no knockdown was detected. P values are from unpaired t-tests. (**F**) Specific ERAP1 inhibitor, ERAP1-IN-1 (CAS: 865273-97-8, MedChemExpress) demonstrates dose-dependent reduction in the percentage of IAV infected cells at 24 hours in A549s. Replicates from 3 separate experiments were normalized to correct for between experiment variation. P-values are from ordinary one-way ANOVA with Dunnett’s multiple comparisons test using DMSO (vehicle) control. (**G**) Interaction between ERAP1 residue 346 and an ERAP1-peptide inhibitor. Ribbon diagram of the crystal structure of ERAP1 (cyan) bound to a 10-mer peptide inhibitor (orange) (PDB:6RQX). ERAP1 residue 346 and the 6^th^ position of the peptide are shown as atom-colored pink and orange spheres, respectively. **(H)** Wild-type ERAP1 G346 and F6 of the 10-mer peptide inhibitor have multiple favorable side chain-backbone contacts. **(I)** However, steric clash between ERAP1 and the 10-mer peptide inhibitor occurs when G346 is mutated to an aspartate using *in silico* mutagenesis. **(J)** Rare rotamers of ERAP1 G346D do not clash with a peptide in which F6 is replace by a glycine but provide no favorable contacts. **(K)** The addition of only a β carbon at position 6 of the 10-mer peptide inhibitor clashes with ERAP1. (**L**) Overexpression of alternative alleles of *ERAP1* in A549 cells demonstrates that the T allele of rs27895 (aspartate) increases the percentage of IAV infected cells at 24 hours. 8 biological replicates from 3 separate experiments were normalized to correct for between-experiment variation. P-values are from ordinary one-way ANOVA with Tukey’s multiple comparisons test. (**M**) Model of how rs27895 affects *ERAP1* transcript and protein and the resulting effect on viral burden. The reference allele of rs27895 encodes cytosine on the genome reference strand and guanine on the transcribed strand. This reference allele encodes glycine at position 346 of ERAP1 and is associated with reduced viral burden in cells. The alternate allele of rs27895 encodes thymine on the genome reference strand and adenine on the transcribed strand. This alternate allele encodes aspartate at position 346 of ERAP1 and is associated with increased viral burden in cells.

While G346D is predicted by SIFT (Sim et al., 2012) to be “tolerated” and by PolyPhen (Adzhubei et al., 2010) to be “benign,” this glycine residue lies within the substrate binding pocket and in the presence of a peptidomimic inhibitor interacts with the first phenylalanine (F6) side chain of this ligand (Giastas et al., 2019) (**Fig. 3G, H**). Moreover, residue 346 is proximal to the ERAP1 active site, which is composed of a catalytic zinc that is coordinated by residues H353, H357 and E376, the proton acceptor residue E354, and “sink” residue Tyr438 (Giastas et al., 2019; Kochan et al., 2011; Nguyen et al., 2011). On the basis of our *in silico* mutational analysis of the structure of the ERAP1-inhibitor peptide complex (Giastas et al., 2019), this SNP encoding a glycine to aspartate substitution, would clearly disrupt the substrate binding pocket of ERAP1 (**Fig. 3I**). Indeed, the carboxylate side chain of an aspartate at ERAP1 residue 346 would result in steric clash with the side chain of any substrate other than those which have a glycine at position 6, including alanine, unless a different route to the active site is taken (**Fig. 3J, K**). Consequently, there would be a significant impact on catalysis or substrate recognition or both. Hence, a G346D substitution in ERAP1 likely has a dual biochemical effect.

Therefore, we tested the effects of overexpression of alternative alleles of *ERAP1* in A549 lung epithelial cells. Strikingly, overexpression specifically of the derived T allele (aspartate; associated with higher IAV burden in scHi-HOST) caused higher IAV burden (p = 0.006), while overexpression of the ancestral C allele (glycine) had no effect (p = 0.7) (**Fig. 3L**). This data confirms the proviral role indicated by the RNAi and inhibitor results and further demonstrates the functional effect of the G346D mutation (**Fig. 3M**).

### Human IAV challenge supports the importance of rs27895 during infection of humans

To determine whether rs27895 was also relevant in humans infected with IAV, we turned to a human IAV challenge study. In the Prometheus study (Grzesiak et al., 2021), volunteers aged 18-55 years were enrolled if baseline antibody titers to the CA09 (influenza A/California/04/09 (H1N1)) strain by hemagglutination inhibition assay were ≤1:10, they were healthy with no co-morbidities or risk factors for severe influenza, and they had no evidence of recent respiratory infection or significant smoking history. Thirty-eight volunteers were inoculated intranasally with CA09 IAV on day 0 and underwent daily assessment and sampling during the subsequent 10 day quarantine period. Modified Jackson symptomology scores were calculated using self-reported symptom diaries (Jackson et al., 1958) and nasal lavage was obtained for qPCR quantification to assess viral burden throughout this period and at follow-up visits up to 6 months post-inoculation. (**Fig. 4A; Table S4**). Both viral burden (by qPCR) and symptoms increased over time among individuals with one copy of the T allele relative to individuals who were homozygous for the C allele. This was confirmed with association testing using EMMAX, controlling for sex, dose, and relatedness of individuals: the T allele of rs27895 was associated with higher qPCR-measured IAV burden at day 4 (p = 0.01; **Fig. 4B; Table S4**) and more severe symptoms from days 3-7, with the most significant association at day 6 (p = 6 x 10^-5^) (**Fig. 4C; Table S4**). These results demonstrate that the association of the rs27895 T allele with higher IAV burden in scHi-HOST (and validated by expression of alternative alleles in A549 cells) is also observed in nasal lavage following experimental human IAV infection, where the T allele correlates with higher viral burden and more severe clinical disease.

**Figure 4.**
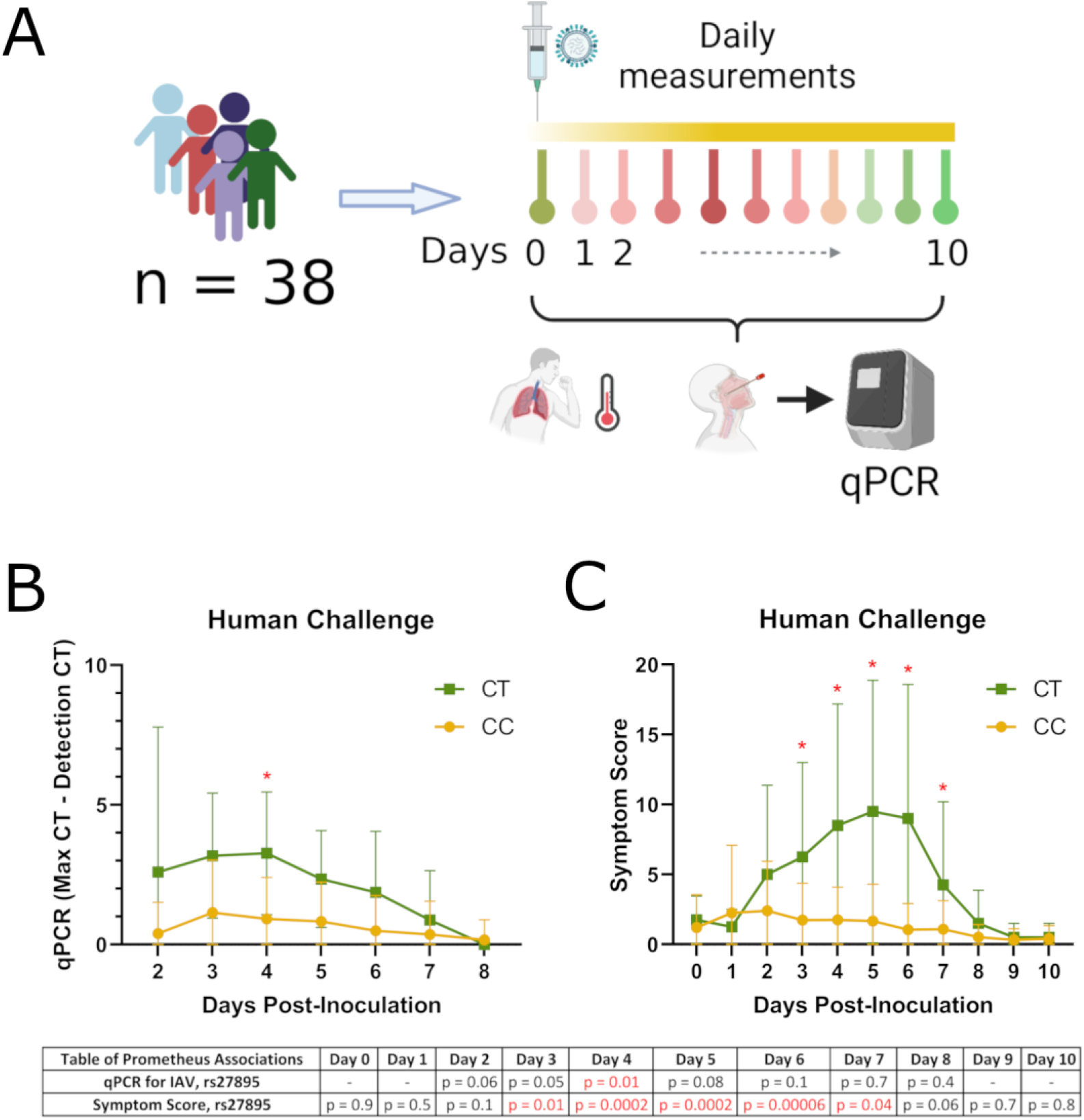
A nonsynonymous variant in *ERAP1* regulates IAV burden and symptomology in human challenge. **(A)** Flow chart of Prometheus study. (**B**) rs27895 T allele is associated with IAV burden at day 4 of the Prometheus Study. Viral burden was measured by nasal lavage by qPCR using pan IAV M gene primers (see Methods). P values from EMMAX at each timepoint for rs27895 are listed below the plot. Mean and standard deviation are plotted with lines connecting means. (**C**) rs27895 T allele is associated with greater symptomology at days 3-7. Jackson symptomology score was assessed daily. P values from EMMAX at each timepoint for rs27895 are listed below the plot. Mean and standard deviation are plotted with lines connecting means.

### Evidence that rs27895 contributes to population differentiation of IAV resistance

IAV has been responsible for at least four pandemics since the start of the 20^th^ century: the swine flu pandemic of 2009-2010, Hong Kong flu of 1968-1970, Asian flu of 1957-1958, and the Spanish flu of 1918-1920 (Saunders-Hastings and Krewski, 2016). The most devastating of these was the Spanish flu, estimated to have resulted in 50 million deaths, or about 5% of the world’s population (Johnson and Mueller, 2002; Taubenberger et al., 2019). Extending further back in history reveals convincing evidence of IAV pandemics since at least the 16^th^ century (Morens et al., 2010). Thus, IAV has had repeated impacts on human populations, consistent with pathogens serving as strong agents of natural selection (Fumagalli et al., 2011; Nedelec et al., 2016; Pittman et al., 2016).

The age and geographic pattern of rs27895 support a model where this SNP was present prior to the out-of-Africa expansion, spread throughout the world, but the resistant, ancestral C allele was selected for by IAV or other pathogens, particularly in East Asian populations. First, rs27895 has an ancient origin based on the Genealogical Estimation of Variant Age approach and resource (Albers and McVean, 2020) (**Fig. S5**). These data indicate the SNP is more than 500,000 years old (assuming a generation time of 25 years), well before the earliest dispersals of *Homo sapiens* out of Africa around 210,000 years ago (Harvati et al., 2019) and consistent with selection at this locus acting on standing variation. Second, the derived T allele of rs27895 is the minor allele throughout the world (MAF = 10% in 1000 Genomes) and is most common in African populations (up to 27% in Mende in Sierra Leonne) (**Fig. 5A**; map generated from (Marcus and Novembre, 2017)). In contrast, the ancestral C allele, associated with IAV resistance, is most common in Asia and has become nearly fixed in East Asian populations (out of 1008 rs27895 alleles, only one is T in CDX, CHB, CHS, JPT, and KHV 1000 Genomes populations). This is confirmed in gnomAD (Karczewski et al., 2020) where only 15 T alleles were detected out of 19,952 East Asian alleles). Notably, for the 10 known IAV pandemics prior to the 2009 swine flu pandemic, all were likely to have originated in Asia (Potter, 2001). This history of emergence is likely secondary to the domestication of ducks and pigs in these regions and their close proximity to humans, which facilitates IAV reassortment and spillover to humans (Saunders-Hastings and Krewski, 2016). Prior to the extensive human migrations that facilitate spread during pandemics, such IAV outbreaks would have been localized to these regions of IAV emergence, acting as a naturally selective force only in these populations to increase the frequency of resistance alleles. Thus, while we cannot be certain that IAV drove the IAV-resistant rs27895 allele to greater frequency in East Asian populations, the combination of genetic association and functional evidence and the likely origin of historical IAV pandemics supports this model.

**Figure 5.**
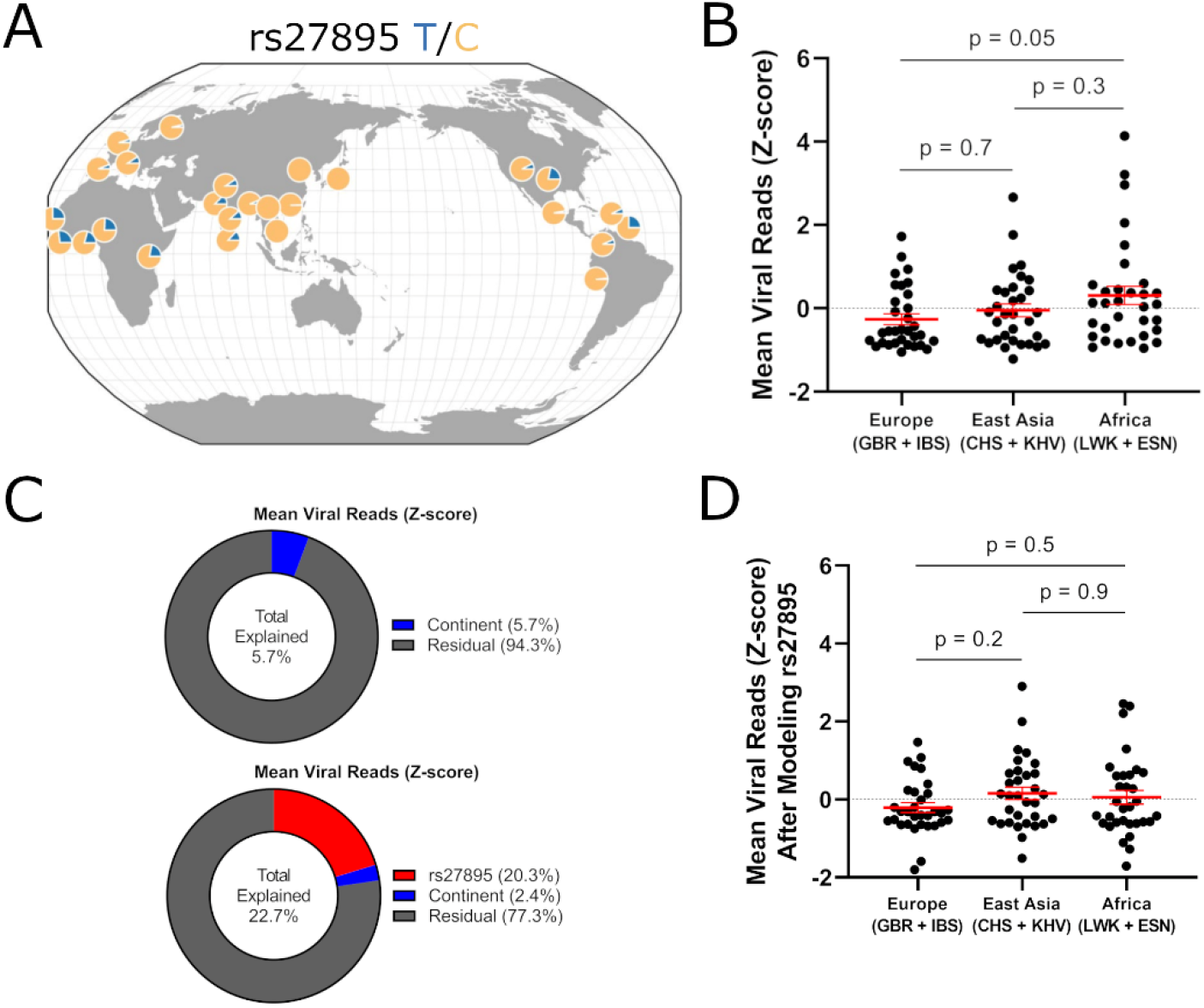
Evidence that population differentiation of rs27895 contributes to genetic resistance to IAV in European and East Asian populations. (A) Geographic distribution of rs27895 is consistent with positive selection of the ancestral C allele in East Asian and European populations. (B) Combined analysis of scHH-LGC and scHH-EIK shows LCLs derived from two European and two East Asian populations are resistant to IAV compared to LCLs from two African populations. P-value from ordinary one-way ANOVA with Tukey’s multiple comparisons test. (C) Linear modeling of the effects of rs27895 and continent on mean IAV burden in LCLs suggests rs27895 contributes to the population differentiation of this phenotype. (Top) Percent explained for continent alone was modeled by lm(Mean Viral Reads ∼ continent). (Bottom) Percent explained for bivariate analysis was modeled by lm(Mean Viral Reads ∼ rs27895 + continent) with continent being modeled on the residual of mean viral reads after regressing out the effect of rs27895. (D) Residual phenotypic variation after removing the effect of rs27895 confirms decrease in total continental effect consistent with known allele frequency differences between populations. P-value from ordinary one-way ANOVA with Tukey’s multiple comparisons test.

If the rs27895 C allele was selected for via pressures from IAV, one would predict greater IAV resistance in regions where the C allele is prevalent. Indeed, we find that LCLs from populations with higher rates of the C allele are overall protected from IAV *in vitro*. A combined dataset of scHH-LGC and scHH-EIK demonstrated lower mean viral reads in LCLs from European populations (GBR+IBS; 91% C allele) and East Asian populations (CHS+KHV; 98% C allele) compared to African populations (LWK+ESN; 75% C allele) (**Fig. 5B**), though the differences within a continental group were much larger than differences between continental groups (**Fig. 5C**, top, continent effect 5.7% of total variance). Linear regression of the effect of rs27895 and continental group on IAV burden demonstrated that the amount of variation explained by continental group is reduced if rs27895 is incorporated into the model (**Fig. 5C**, bottom, rs27895 effect 20.3% total variance, continental effect 2.4% of residual variance). Specifically, removing the effect of rs27895 reduces IAV burden in the African continental group (median from 0.11 to - 0.21), consistent with the higher frequency of the susceptible derived allele in Africa. Additionally, removing the effect of rs27895 increased residual burden in the European continental group (median from -0.55 to -0.33) and East Asian continental group (median from -0.14 to 0.10), consistent with the low frequency of the susceptible derived allele in Europe and even lower frequency in East Asia (**Fig. 5D**, compare to **Fig. 5B**).

## Discussion

Influenza has been one of the largest causes of human suffering and mortality in modern times. Since 1918, global IAV infections have been a yearly occurrence, with ∼30 million symptomatic infections in the United States alone (CDC, 2020). However, there are broad differences in influenza susceptibility and severity, with outcomes from asymptomatic infections (∼16%(Leung et al., 2015)) to death (0.2% in 2014-2015(CDC, 2020)). Here we have developed a rapid scRNA-seq method to identify genetic susceptibility to IAV cellular infection. Integration with human IAV challenge data revealed human genetic differences that confer susceptibility to IAV infection in cells and people.

The scHi-HOST method has advantages of speed and scalability. Once an infection assay has been optimized for a particular pathogen, scHi-HOST can be carried out and analyzed within a few weeks. However, this requires that LCLs can serve as a relevant model for cellular infection and optimization of timing and dose; utility is limited in cases where the receptor in not well expressed in LCLs. In the case of IAV, LCLs are highly infectable, consistent with B cells being direct targets for IAV entry (Dougan et al., 2013; Lersritwimanmaen et al., 2015; Tumpey et al., 2000), proliferating in response to IAV (Anders et al., 1984; Poumbourios et al., 1987; Rott and Cash, 1994) and eventually undergoing cell death (Nichols et al., 2001; Tumpey et al., 2000), likely contributing to lymphopenia during severe infection (Shinde et al., 2011). Additionally, IAV infection induces global transcriptional changes in B cells, many mediated by IFN (Chang et al., 2007; Priest and Baumgarth, 2013). As new IAV threats emerge from animal reservoirs, testing these new reassorted strains with scHi-HOST could serve as an early warning of pandemic potential as well as common genetic resistance to these new strains in human populations. This application of scHi-HOST may be particularly relevant with the Centers for Disease Control and Prevention tracking and evaluating potential for emergence and public health impact of IAV strains currently circulating in animals but not humans (Cox et al., 2014). Testing such strains with scHi-HOST could further evaluate infection potential across diverse humans while also revealing possible resistance alleles against these emerging strains. scHi-HOST can also be increased in scale for greater power. Our proof-of-concept using 96 LCLs allowed for hundreds of individual cells to be assayed for each LCL but required narrowing the genetic search space to nonsynonymous variants with high minor allele frequencies to reveal the SNP in *ERAP1* that was validated experimentally and in human challenge volunteers. Notably, the genome-wide significant hit identified by scHi-HOST (rs1923314) does show an association with the COVID19hg A1 phenotype (very severe respiratory confirmed covid vs. not hospitalized covid; p = 0.0004; COVID-19 HGI data freeze 4; (Initiative, 2020)) with directionality consistent with scHi-HOST (A allele associated with severe COVID and greater mean IAV burden), but further work is necessary to validate the relevance of this SNP to viral infection. As scRNA-seq throughput and cost continue to improve, we anticipate increasing the cells in a single scHi-HOST experiment by an order of magnitude or more, likely revealing high confidence genome-wide significant hits for further validation experimentally and in human challenge datasets. Finally, while the variants identified by scHi-HOST are currently restricted to those with cell autonomous effects (or at least those that require only interactions of a single cell type), future developments may incorporate expansion of the method and computational pipeline to other cell types or even complex organoid models. In this regard, while *TNFSF12* did not have a phenotype by RNAi in LCLs, the secreted product of this gene (and others implicated by eQTLs in our analysis) may have important non-cell-autonomous effects in IAV infection that require more complex models to detect.

The simultaneous identification of eQTLs and SNPs associated with more complex cellular phenotypes in scHi-HOST allows for integrated identification of SNPs whose effects on gene expression help dictate the outcome of cellular host-pathogen interactions. Most eQTLs and even response eQTLs did not contribute to control of IAV burden in LCLs. This is a similar challenge to immunologists who confront the fact that ISGs comprise perhaps 10% of the genome, but individually most have little effect against an individual infection (Schneider et al., 2014; Schoggins, 2019). Regardless, our finding that high expression of IFNs and ISGs across LCLs prior to infection is correlated with lower IAV burden confirms the importance of these genes in aggregate and warrants additional experimental dissection. However, our identification and experimental validation of a nonsynonymous variant in *ERAP1* as a major regulator of IAV in cells and humans underscores that despite noncoding variants receiving much attention as accounting for most of the identified GWAS risk alleles of common diseases (Maurano et al., 2012), coding variation should not be ignored in trying to understand human genetic susceptibility to infectious diseases.

How ERAP1 serves as a proviral factor in IAV infection is unknown. The canonical function of ERAP1 is to trim proteosome-generated peptides from the N-terminus to a preferred 9 residue length for MHC Class I loading. MHC I is then trafficked from the endoplasmic reticulum to the cell surface for surveillance by T-cell receptors on cytotoxic T cells (Saulle et al., 2020). This has been reported to be an IAV-regulated process: infection with H1N1 activates p53 to increase *ERAP1* expression, resulting in increased MHC I presentation (Wang et al., 2013). Mouse ERAP trims an immunodominant IAV epitope (nucleoprotein (NP) 383-391) for presentation by human HLA-B27 (Akram et al., 2014). This same study reported that B27 transgenic/*ERAP^-/-^* mice have greater edema and weight loss during IAV infection, suggesting an antiviral role in mice. Our data indicate that ERAP1’s role may be complex, with our association and functional data demonstrating a proviral role for ERAP1, dependent on the rs27895 G346D polymorphism. Several other ERAP1 missense variants have been identified as important for disease, including rs30187 (K528R) associated with ankylosing spondylitis (AS; p = 4 x 10^-45^; (International Genetics of Ankylosing Spondylitis et al., 2013) and rs27044 (Q730E) associated with psoriasis (p = 8 x 10^-21^; (Zuo et al., 2015)). These SNPs do not show an association with mean IAV burden in scHi-HOST (p = 0.73 and p = 0.57). Conversely, rs27895 is not associated with ankylosing spondylitis (p = 0.83 from downloaded summary statistics from EBI-GWAS catalog; (International Genetics of Ankylosing Spondylitis et al., 2013)) or psoriasis (p=0.40 from downloaded summary statistics from EBI-GWAS catalog; (Tsoi et al., 2012)). This specificity highlights that different ERAP1 variants have different roles in diseases that may be due to specific changes in the MHC I antigen repertoire, and that missense mutations likely have different effects on IAV infection than complete loss-of-function. How changing the composition of the MHC I antigen repertoire (or the peptide composition within the ER lumen) could impact IAV infection is unclear, and while others have examined how the *ERAP1* haplotype associated with AS impacts the peptidome of presentation by HLA-B*27 (Lopez de Castro, 2018), consequences of the haplotype containing rs27895 (Hap5 from (Ombrello et al., 2015)) has been largely unexplored.

Alternatively to affecting MHC-I presentation, ERAP1 has functions beyond MHC I peptide trimming that may contribute to its role in IAV infection. This includes regulation of immune cell activation and cytokine production through secreted ERAP1 (Aldhamen et al., 2015; Tsujimoto et al., 2020). However, scHi-HOST is a pooled infection assay, and thus would not be expected to identify genetic differences that impact secreted factors acting through non-cell-autonomous effects. ERAP1 also regulates ectodomain shedding of cell surface receptors (Cui et al., 2003a, b). In considering this function, ERAP1 may regulate cleavage of transmembrane receptors that are trafficked through the ER. Thus, while our work demonstrates the importance of rs27895 on IAV infection in both cells and humans, future studies will decipher how ERAP1 D346 specifically exerts its proviral function.

While the geographic distribution of rs27895 considering the history of IAV epidemics and our association and functional data suggests variation in *ERAP1* may have helped humans adapt to IAV infection, human evolution of complex cellular phenotypes of infection are undoubtedly polygenic. Even within the *ERAP1* gene, the different variants associated with disease likely have been driven by human pathogens that have varied in time and space. Our data with LCLs showing IAV resistance in broadly European and East Asian populations contrasts with recent data using monocytes and PBMCs that shows resistance in African populations (O’Neill et al., 2021; Randolph et al., 2021). What may be critical is the timing of measurement, with these studies using an early 6-hour timepoint compared to our 24-hour timepoint. Indeed, Randolph et al. show that greater European ancestry is associated with higher IFN levels and higher IAV burden at 6 hours, but that by 24 hours, PBMCs with a higher IFN response had substantially lower IAV burden (Randolph et al., 2021). This is reminiscent of our finding that IFNα levels prior to infection are correlated with lower burden by 24 hours (see Figure 1K). Differences in cell type may also play a role as Randolph et al. demonstrate many cell-type specific ancestry effects on IAV-induced transcriptional changes, though they also note that the ancestry effect on the IFN response is broadly conserved across all PBMC cell types.

Ultimately, we hope for broad adoption of scHi-HOST as a generalizable tool for rapid assessment of genetic susceptibility and resistance to pathogens that are important for human health or that pose an emerging threat. We have developed scHi-HOST to be a truly accessible resource; scHH-LGC and scHH-GBK aliquots are in liquid nitrogen storage, to be shared with any labs that have developed an infection phenotype in LCLs. In addition to surveillance of emerging IAV strains, scHi-HOST can readily be applied to other pathogens with polyadenylated transcripts for identifying eQTLs in response to pathogen, population differences in resistance and susceptibility, and the genetic differences underlying this variation. Such genetic differences could reveal targets for drug development or FDA-approved drugs for repurposing as important prophylactics or therapeutics for future pandemics.

## Methods

### Lymphoblastoid cell lines

1000 Genomes LCLs from LWK (Luhya in Webuye, Kenya), ESN (Esan in Nigeria), GBR (British in England and Scotland), IBS (Iberian in Spain), CHS (Han Chinese South), and KHV (Kinh in Ho Chi Minh City, Vietnam) populations were purchased from the Coriell Institute. LCLs were maintained at 37°C in a 5% CO2 atmosphere and were grown in RPMI 1640 media (Invitrogen) supplemented with 10% fetal bovine serum (FBS), 2 mM glutamine, 100 U/ml penicillin-G, and 100 mg/ml streptomycin.

### Viruses

The PR8 (A/Puerto Rico/8/1934) mNeon-HA virus was generated and validated as previously described (Harding et al., 2017). Both the wild-type and PR8 mNeon-HA viruses were amplified by via injection of the respective virus into 10-day-old embryonated hen eggs (Charles River) for three days at 37C. After incubation, allantoic fluid containing the virus was harvested, briefly spun down to remove debris, and frozen at -80°C. Viruses were then titered, as previously described, via standard plaque assay on Madin-Darby canine kidney cells (MDCKs). For the human challenge study, a live wild-type GMP-certified influenza A/California/04/2009 (H1N1) was a gift from Altimmune and produced using standard methods certified by the company.

### Viral Infection of pooled lymphoblastoid cells

48 LCLs were pooled in equal numbers and added to a 24-well plate in PBS with 0.35% BSA, 2 mM glutamine, 100 U/ml penicillin-G, and 100 mg/ml streptomycin. Cells were infected with A/Puerto Rico/8/1934 at MOI 50 or left as uninfected controls. At 3 hours post infection, each well was spiked with 600 µl of RPMI 1640 media (Invitrogen) supplemented with 10% fetal bovine serum (FBS), 2 mM glutamine, 100 U/ml penicillin-G, and 100 mg/ml streptomycin. At 24 hours post infection, cells from each sample were collected, spun down, and resuspended in PBS with 0.04% BSA for single-cell cDNA library preparation. LCLs from LWK, GBR, and CHS were used in scHi-HOST-LGC. LCLs from ESN, IBS, and KHV were used in scHi-HOST-EIK.

### Single-cell RNA-seq cDNA library preparation

Cell samples were counted and checked for viability on a Guava EasyCyte HT system by 7-AAD staining before they were diluted to 1 million cells/mL with an intended capture of 10,000 cells/sample. Each individual sample was used to generate individually barcoded cDNA libraries using the 10x Chromium Single Cell 3’ platform version 3.1 (Pleasanton, CA) following the manufacturer’s protocol. The Chromium Controller partitions the cells into nanoliter-scale gel beads in emulsion (GEMS) within which cell-specific barcoding and oligo-dT-primed reverse-transcription occurs.

### Sequencing of single-cell cDNA library

cDNA samples from scHi-HOST-LGC were dual-indexed and sequenced on one Illumina NovaSeq S4 flow cell with target depth 100,000 reads per barcoded droplet. Reads were sequenced with read 1 length of 28 base pairs (bp) and read 2 length of 150 bp. cDNA samples from scHi-HOST-EIK were single-indexed and sequenced on an Illumina HiSeq system with a target depth of 50,000 reads per barcoded droplet. Reads were sequenced with read 1 length of 150 bp and read 2 length of 150 bp.

### Single-cell RNA-seq alignment

Raw sequencing results were processed using the 10X Genomics CellRanger 4.0. Reads from each sample were mapped to GRCh37. Reads from infected samples were also mapped to the A/Puerto Rico/8/1934 genome. For scHi-HOST-EIK, since the library was single-indexed, we used 10X Genomics Index-hopping-filter to remove index-hopped reads (https://github.com/10XGenomics/index_hopping_filter) before mapping to either human or viral genome.

### Assignment of reads to each of 48 LCLs

To assign each barcoded read to an LCL identity, we used Demuxlet (Kang et al., 2017). Demuxlet takes a bam file with barcoded sample reads and a VCF (obtained from the 1000 Genomes Project, phase 3 release) to computationally determine the LCL contained in each GEM reaction. With barcodes assigned to each of 48 LCLs, we used Subset-Bam (10X Genomics) to subset reads from each sample alignment bam into 48 LCL-specific bam alignment files. From these LCL-specific alignment bams, we used HTSeq-Count (Anders et al., 2019) to produce counts files for differential expression, eQTL discovery, and other transcriptomic analyses.

### Single cell RNA-seq analysis using Seurat

With the filtered and annotated expression matrix outputs of the CellRanger 4.0 pipeline, we used Seurat V4 (Hao et al., 2021) to perform single-cell analysis of PR8-infected and uninfected LCLs. Seurat data was annotated with LCL identity and normalized viral reads per droplet for analysis. Analysis on feature counts data was performed using standard log-normalization. Droplets with > 20% mitochondrial reads were excluded from analysis. UMAP clustering of combined scHi-HOST-LGC and scHi-HOST-EIK datasets was performed with 15 dimensions and resolution of 1.2.

### Differential gene expression using HTSeq-Count and Deseq2

After subsetting sample alignments to LCL-specific alignments, feature counting was performed using HTSeq-Count (Anders et al., 2019) with Gencode v19 feature coordinates. All differential expression testing was performed in R using the DESeq2 (Love et al., 2014) package. Normalization of the raw counts matrix was achieved using DESeq2’s default “Median of Ratios” method. Briefly, DESeq2 computes a pseudo-reference sample using the geometric mean for each gene. It then calculates a ratio of each sample to the computed pseudo-reference for each gene. The normalization factor (“size factor” in DESeq2) for each sample is computed by taking the median of the ratios of each gene to the pseudo-reference for each sample. Finally dividing the raw counts by the size factor yields normalized counts for further analysis in DESeq2. PCA of uninfected and infected cells was performed using variance-stabilizing transformation (vst) of the counts matrix and principal components were calculated on the 40,000 most variably expressed features.

### Allele specific expression analysis using RASQUAL

We used RASQUAL v1.1 (Kumasaka et al. 2016) to identify allele-specific eQTLs (aseQTLs). RASQUAL uses a probability decomposition approach to estimate aseQTLs by jointly modeling both total allele-specific count and total fragment count. RASQUAL requires two main input files, a genotype file (vcf format) including the phased genotypes and reads counts for reference allele and alternative allele for each variant for each sample, and a gene expression file including read counts of all genes.

We downloaded phased genotype data (vcf format in GRCh37) from the 1000 Genomes Project Phase 3 (available at http://ftp.1000genomes.ebi.ac.uk/vol1/ftp/), and filtered using bcftools v1.9 with parameters of “-q 0.01:minor -m 2 -M 2” to keep only biallelic SNPs and those with minor allele frequency more than 0.01. LCL-specific transcriptome alignments were obtained from Demuxlet and Subset-Bam as above. Given phased genotypes and individual sample alignment, we then quantified allele-specific read counts overlapping with biallelic variants using the ASEReadCounter function from GATK v3.8. The following parameters were used: “-U ALLOW_N_CIGAR_READS -minDepth 1 --allow_potentially_misencoded_quality_scores -- minMappingQuality 10 --minBaseQuality 10”.

For gene expression input files, featureCounts (Liao et al., 2014) was used to count reads per gene for each individual against GRCh37 reference genome with default parameters. We filtered out low expression and variable genes with < 5 reads in < 5 LCLs. Gene read counts were then corrected using RASQUAL’s GC-correction to remove potential sample-specific GC bias. We included sex, top 2 genotypic PCs and transcriptome library size as covariates calculated using Plink v1.9 (Purcell et al., 2007). For each gene, SNPs within a 1 megabase window from each gene’s start and end position were included in our analysis. The final command and parameters were “tabix filtered.genotype.vcf.gz Chromosome:regionStart-regionEnd | rasqual -y expression_Y.bin -k size_factor.K.bin -n NumberSampleSize -l NumberCisSNPs -m NumberFeatureSNPs -s exon_startposition -e exon_endposition -t -f GeneName”.

RASQUAL’s “--random-permutation” option was performed to calculate the permuted p values and construct the empirical null distribution. The nominal p values of all variants within each gene were corrected using the Benjamini-Hochberg method, and significant aseQTLs for each gene was defined using a 5% false discovery rate (FDR).

### Viral Burden Phenotyping

With human and viral reads assigned to each droplet, we calculated viral burden for each droplet as a function of the total reads mapped to that droplet to control for differences in read depth between droplets, resulting in normalized counts of viral reads per droplet. To get burden per LCL phenotypes, we used Demuxlet assignments of droplets to LCLs to create a distribution of viral burden for each LCL. We used the mean of the distributions as phenotype for GWAS.

### GWAS of viral burden

GWAS analysis on viral burden as calculated above was performed using a linear mixed model implemented in EMMAX with default parameters (Kang et al. 2010). The input genotypes for each LCL were obtained from the 1000 Genomes Project described as above. EMMAX controls for population stratification and cryptic genetic relatedness by incorporating a kinship matrix. The “emmax-kin” function was used to calculate the pairwise genetic relationships among individuals with Balding-Nichols algorithm. We also incorporated sex to control potential sex influence on phenotypes. In this analysis, we only carried out analysis for SNPs with minor allele frequency > 10%. Furthermore, we excluded all SNPs from analysis significantly deviating from Hardy-Weinberg equilibrium (p < 10^-4^) in any of the 6 populations to remove variants that may have been affected by genotyping error. After filtering for minor allele frequency and Hardy-Weinberg equilibrium we are left with 5.2 million SNPs for GWAS. P-values reported are corrected for genomic inflation factor (λ = 1.05).

### Enrichment analysis of eGenes

The enrichment of scHi-HOST eGenes in GTEx and enrichment of ISGs in scHi-HOST eGenes were tested using Fisher’s exact test. Significance of enrichment was calculated using the following equation:

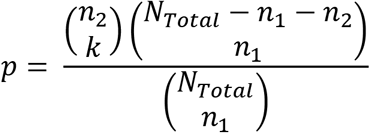

Where *N_Total_* is the total number of genes, the *n*_1_ and *n*_2_ are the number of eGenes associated with scHi-HOST and GTEx, and *k* is the number of eGenes shared between scHi-HOST and GTEx.

### Viral infection of human volunteers

The Prometheus human infection challenge study of healthy adult volunteers with Influenza A/California/04/09 (H1N1-2009) was previously described (Grzesiak et al., 2021). This study was performed at Imperial College London (London, UK) in accordance with the protocol, the Consensus ethical principles derived from international guidelines including the Declaration of Helsinki and Council for International Organizations of Medical Sciences (CIOMS) International Ethical Guidelines, applicable ICH Good Clinical Practice guidelines, and applicable laws and regulations. IAV challenge study was reviewed and approved by the Institutional Review Board at Duke University and the UK Health Research Authority London-Fulham Research Ethics Committee (Ref. 17/LO/0965). Written informed consent was obtained from all participants, who provided informed consent for all study procedures including genetics, before screening and enrollment. Each volunteer was pre-screened for serum antibodies against the CA09 strain by hemagglutination inhibition, followed by screening of risk factors for severe disease and at-risk close contacts. Individuals were enrolled if HI antibody titers against the inoculating influenza strain were ≤1:10. Exclusion criteria included current pregnancy, chronic lung disease, smoking, bronchodilator or steroid use (last 12 months), allergic symptoms or medication for allergic symptoms (last 6 months), acute respiratory infection (last 6 weeks), or immunodeficiency. All participants were screened for further exclusion due to drug use, pregnancy, and allergic reactions that would preclude vaccine use. Screening tests included a panel of routine blood tests, tests for immune deficiency and blood-borne virus serology. Subjects were inoculated intranasally with live GMP-certified influenza A/California/04/2009 (Altimmune) at a mean dose of 10^6^ within 6 weeks of screening. Post-inoculation, subjects were confined to an isolation facility for eight days with daily collection of clinical and physiologic findings, modified Jackson symptomology scores (Jackson et al., 1958), nasal lavage, and blood samples. Symptom scores were recorded up to day 10 post-inoculation and follow-up interviews and physical examinations were conducted at day 14 and 28.

### Quantitative PCR of nasal lavage fluid

Quantitative PCR was performed on nasal lavage samples as previously described (Jozwik et al., 2015). Briefly, total RNA was isolated using the QIAamp Viral RNA kit (Qiagen) according to the manufacturer’s instructions. Reverse transcription of 13 µl of total isolated RNA was achieved using the High Capacity RNA-to-cDNA kit (Applied Biosystems) according to the manufacturer’s instructions. Quantitative RT-PCR reactions for viral load were achieved using pan-IAV M gene primers and probes (forward: GACCRATCCTGTCACCTCTGAC, reverse: AGGGCATTYTGGACAAAKCGTCTA, probe: TGCAGTCCTCGCTCACTGGGCACG) with the TaqMan Universal Master Mix II (Applied Biosystems) and 7500 Fast Real-Time PCR System (Applied Biosystems). Absolute quantification was calculated using a plasmid DNA standard curve.

### Prometheus low-pass whole genome sequencing and imputation

Whole genome sequencing was performed on Prometheus human challenge study subjects. Total genomic DNA was extracted from buffy coat using the QIAGEN DNeasy blood kit following manufacturer’s instructions (average genomic DNA concertation from 38 volunteers was 95.58 ng/µl). Whole genome sequencing was carried out by BGI using BGI DNBSEQ low-pass genome sequencing to 4x coverage. Short reads were aligned to human genome GRCh37 and imputed through the Gencove ImputeSeq pipeline using the 1000 Genome Project Phase 3 reference panel (Li et al., 2021). Imputed variants were filtered using *bcftools* v1.9 (Danecek et al., 2021), including minor allele frequency > 0.05, genotype missingness < 0.2, sample missing calls < 0.2. In the subsequent analysis, only autosomal biallelic variants were included. EMMAX was used to run GWAS on qPCR of IAV load and symptom scores, with details described above.

### LCL RNAi Experiments

LCLs (2 x 10^5^ cells) were treated for three days in 500 μL of Accell media (Dharmacon) with either non-targeting Accell siRNA #1 or an Accell SmartPool directed against human *ERAP1* or *TNFSF12* (1 μM total siRNA; Dharmacon) in a 24-well TC-treated plate. After 3-day incubation with siRNA, cells were plated at 30,000 per well in 50μl PBS with 0.35% BSA, 2 mM glutamine, 100 U/ml penicillin-G, and 100 mg/ml streptomycin in 96-well plates. Infections were conducted at MOI 50 with PR8-mNeon-HA for 3 hours before wells were spiked with 75µl RPMI (10% FBS + 1% Pen-Strep). Cells were assayed for Percent mNeon+ cells using a Guava Easycyte flow cytometer at 24 hours post-infection. RNA was collected from 2 x 10^5^ pelleted cells using the RNeasy mini kit (Qiagen). qPCR using primers specific for *ERAP1* or *TNFSF12* were used to validate knockdown relative to 18S rRNA.

### ERAP1 Inhibitor in A549 Experiments

A549 cells were plated at 25,000 cells per well in 50 µl growth media (DMEM + 10% FBS + 1% Pen-Strep) in 96 well plates. 1 day after plating, media was removed and replaced with 50 µl PBS with 0.35% BSA, 2 mM glutamine, 100 U/ml penicillin-G, and 100 mg/ml streptomycin and 1 µM, 5 µM, or 10 µM ERAP1-IN-1 (CAS: 865273-97-8, MedChemExpress), or equal volume DMSO (vehicle). After 30-minute incubation, cells were infected at MOI 1 with PR8-mNeon-HA for 1 hour before infectious media was removed and replaced with post-infection media (Opti-mem + 0.01% FBS + 1% Pen-Strep + 0.35% BSA). 1-day post-infection, cells were trypsinized and assayed for percent mNeon+ cells using a Guava Easycyte flow cytometer.

### Overexpression in A549

For lentivirus production, HEK-293T cells were transfected with 1 µg pLEX-ERAP1, 0.4 µg pMD2.G and 1 µg pCMVΔ8.74. Lentiviral supernatant media was collected after 72 hours. Next, A549 cells in a 24-well plate were transduced with 1 mL lentivirus and selected with 1 µg/mL puromycin after 48 hours. After 3 days selection, surviving cells were expanded.

Cell lines were plated at 25,000 cells per well in 50 µl selection media (DMEM + 5% FBS + 1% Pen-Strep + 1% Puromycin) in 96-well plates. 1 day after plating, media was removed and replaced with 50 µl PBS with 0.35% BSA, 2 mM glutamine, 100 U/ml penicillin-G, and 100 mg/ml streptomycin and infected at MOI 0.1 with PR8-mNeon-HA for 1hr before infectious media was removed and replaced with selection media. 1 day post-infection, cells were trypsinized and assayed for percent mNeon+ cells using a Guava Easycyte flow cytometer.

### *In silico* mutagenesis of ERAP1 and 10-mer peptide

*In silico* mutational analysis of ERAP1 bound to the 10-mer peptide (PDB: 6RQX) was performed using the Pymol mutagenesis function (The PyMol Molecular Graphics System, Version 1.8.6.0 Schrodinger, LLC).

### Data and Software Availability

RASQUAL and EMMAX results are available for download at the Duke Research Data Repository https://doi.org/10.7924/r4g163n0s

scRNA-seq data will be available in GEO upon publication.

## Acknowledgments

We thank the investigators and individuals from diverse populations genotyped as part of the 1000 Genomes Project, who have made their LCLs available through the Coriell Institute. We thank the human volunteers in the Prometheus IAV challenge study. We thank the members of the Ko lab for useful discussion during the development of scHi-HOST. We would like to acknowledge the assistance of the Duke Molecular Physiology Institute Molecular Genomics Core for the generation of data for the manuscript. We thank Vincent Gardeux for making the low-memory modification to Demuxlet allowing us to demultiplex 48 individuals on a RAM-limited node. BHS, LW, and DCK were supported by NIH R21AI153812. TWB, CC, MTM, and CWW were supported by DARPA grant N66001-17-2-4014 (Prometheus Study). JSB was supported by NIH 1F31AI143147. BHS was supported by a Precision Genomics Collaboratory-OBGE Graduate Student Pilot Research Grant Award. DCK was supported by a Duke University SOM Core Facility Voucher Program. CC was supported by the Biomedical Research Centre (BRC) award to Imperial College Healthcare NHS Trust. Infrastructure support was provided by the NIHR Imperial Biomedical Research Centre and the NIHR Imperial Clinical Research Facility. The views expressed are those of the authors and not necessarily those of the NHS, the NIHR or the Department of Health and Social Care.

## Author Contributions

Conceptualization: BHS, LW, ERK, SG, NSH, DCK; Methodology: BHS, LW; Formal Analysis: BHS, LW; Investigation: BHS, LW, XZ, ATH, JSB, EJW, TWB, JA, EB, ZG, SP, RGB; Resources: ERK, TWB, JA, CC, MTM, CWW, NSH; Data Curation: BHS, LW; Writing – Original Draft: BHS, LW, ERK, DCK. Writing – Review & Editing: All authors; Visualization: BHS, LW, EJW, DCK; Supervision: RGB, CWW, SG, NSH, DCK; Funding Acquisition: CC, CWW, DCK.

## Supplemental Information

**Figure S1. LCL properties that do not correlate with IAV burden.** (**A**) EBV copy number in LCLs does not correlate with IAV burden. EBV copy number from 72 LCLs out of 96 LCLs are obtained from Mandage et al. (2017). A non-significant correlation (p = 0.8) was observed between EBV copy number and mean viral reads (normalized per cell) using Spearman correlation. Each point with distinct color represents an LCL. (**B**) The number of each LCL recovered for RNA-seq does not correlate with IAV burden. A non-significant correlation (p = 0.6) was observed between cell number and mean viral reads (normalized per cell) using Spearman correlation. Each point with distinct color represents an LCL.

**Figure S2. IFNα genes correlate with viral burden.** A) Expression of all detected IFNα genes vs. log2(Mean Normalized Viral Reads) shows that, for expressed IFNα genes, higher basal expression is correlated with lower viral burden after infection. Correlation coefficient and p-values are from Spearman’s correlation. B) Induction (log2(IAV Counts + 1 / Uninfected Counts + 1) of all IFNα genes reveals that IFNα induction is correlated with higher viral burden. Correlation coefficient and p-values are from Spearman’s correlation.

**Figure S3. Cellular GWAS of mean IAV burden in LCLs.** (A) Manhattan plot of mean IAV burden based on p values from EMMAX using kinship matrix, sex as a covariate, and with SNPs with MAF < 10% and deviation from Hardy-Weinberg Equilibrium (p < 1×10-4) excluded. P values plotted are corrected for genomic inflation factor (λ = 1.05). (B) QQ plot of the same data in A.

**Figure S4. Genotype median plot for rs27895 mean IAV reads in individual scHi-HOST populations.** Plotting by continent reveals that rs27895-T is associated with increased viral reads in European and African populations while the C allele of rs27895 is nearly fixed in East Asian populations.

**Figure S5. Estimate of the age of rs27895 suggests that the variant was present in human populations before the out-of-Africa expansion.** Data and figure from https://human.genome.dating

**Table S1. Differentially expressed genes for scHH-LGC comparing uninfected vs. IAV-infected cells.** Normalization and differential expression testing performed using DESeq2.

**Table S2. GSEA for scHH-LGC comparing uninfected vs. IAV-infected cells using the C2 gene sets.** Enrichment for gene sets was calculated using pre-ranked input. Metric for GSEA was sign(Fold-change)*(-log10(adjusted p-value)).

**Table S3. eGenes identified in scHi-HOST using RASQUAL.** For each gene, only the lead SNP was retained, then 2-step adjusted p values were applied to all genes. eGenes were defined as genes with lead SNP 2-step p values < 0.05. Data files for eQTLs are available for download from Duke Research Data Repository https://doi.org/10.7924/r4g163n0s.

**Table S4. Characteristics of the Prometheus IAV Challenge and Association of rs27895.**

