## Supplementary material for "Single-cell genome-wide association reveals a nonsynonymous variant in *ERAP1* confers increased susceptibility to influenza virus": Figures S1-S5

**Supplemental Information**

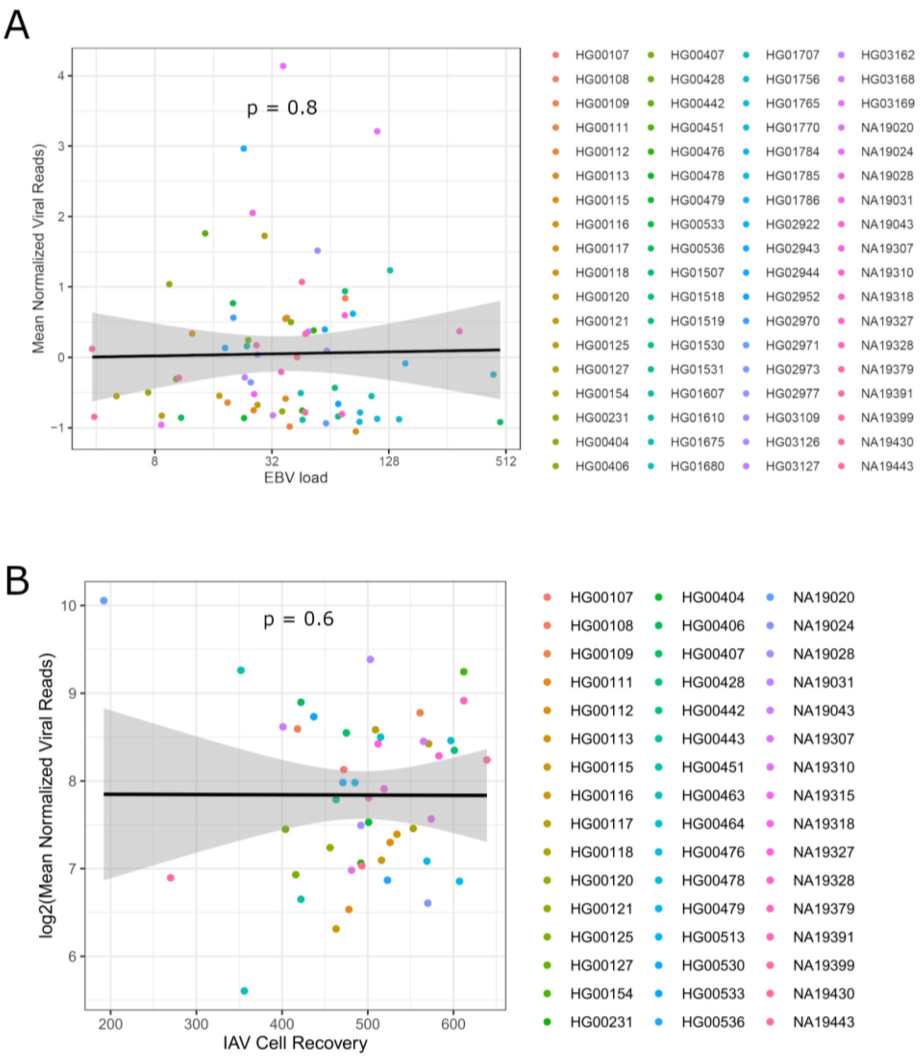


**Figure S1.** LCL properties that do not correlate with IAV burden. (**A**) EBV copy number in LCLs does not correlate with IAV burden. EBV copy number from 72 LCLs out of 96 LCLs are obtained from Mandage et al. (2017). A non-significant correlation (p = 0.8) was identified between EBV copy number and mean viral reads (normalized per cell) using linear regression. Each point with distinct color represents an LCL. (**B**) The number of each LCL recovered for RNA-seq does not correlate with IAV burden. A non-significant correlation (p = 0.6) was identified between cell number and mean viral reads (normalized per cell) using linear regression. Each point with distinct color represents an LCL.

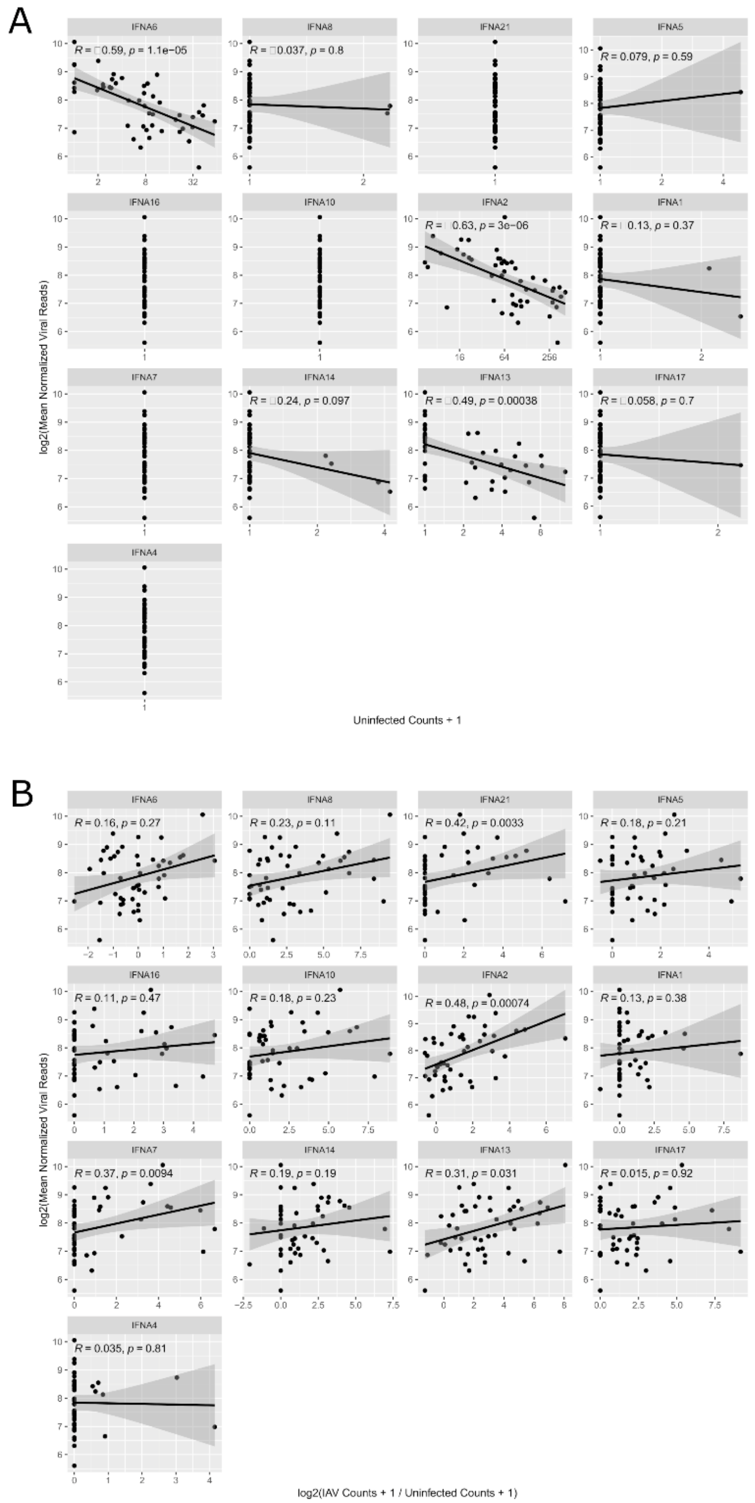


**Figure S2**. **IFNα genes correlate with viral burden.** A) Expression of all detected IFNα genes vs. log2(Mean Normalized Viral Reads) shows that, for expressed IFNα genes, higher basal expression is correlated with lower viral burden after infection. Correlation coefficient and p-values are from Spearman's correlation. B) Induction (log2(IAV Counts + 1 / Uninfected Counts + 1) of all IFNα genes reveals that IFNα induction is correlated with higher viral burden. Correlation coefficient and p-values are from Spearman's correlation.

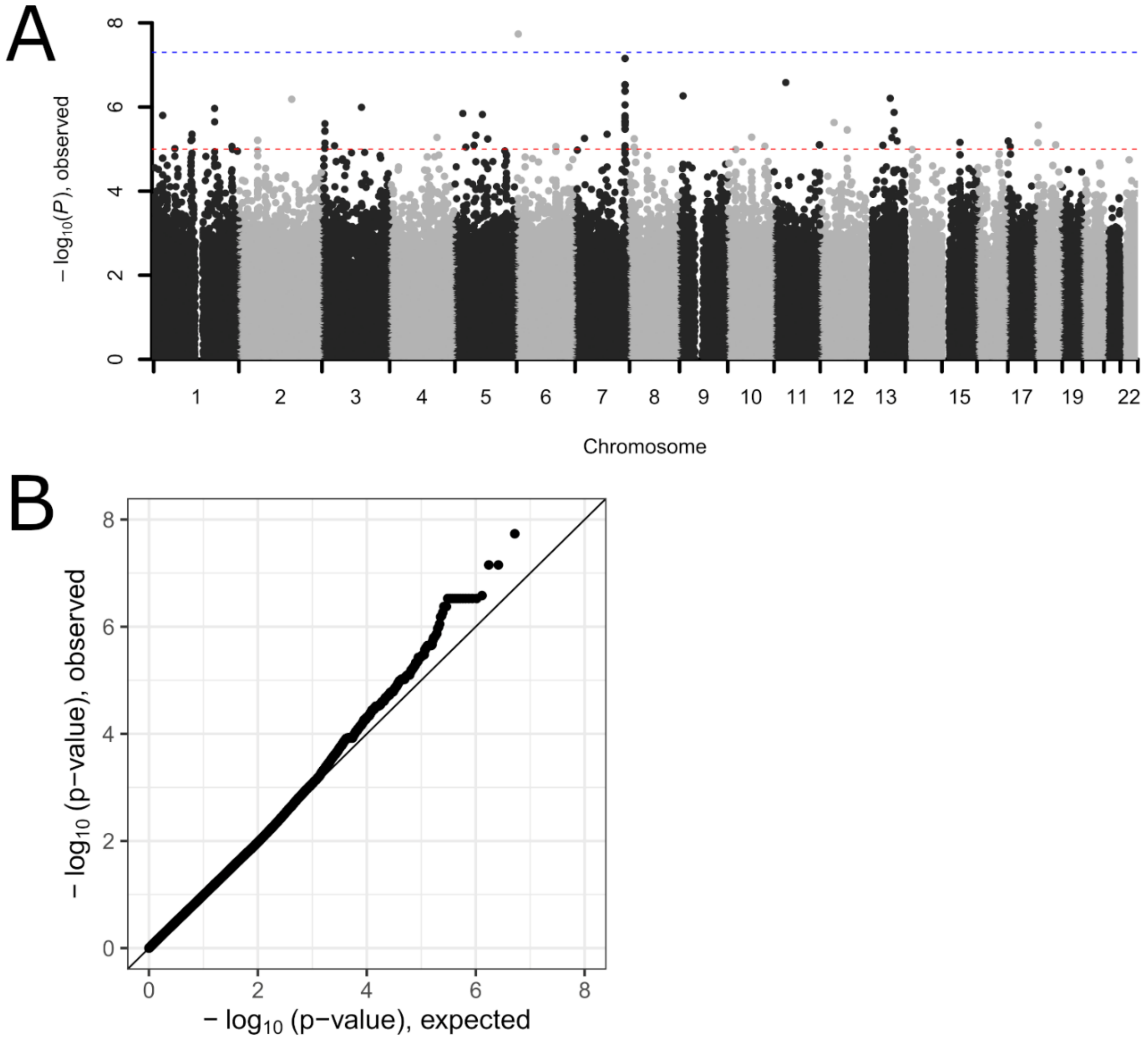


**Figure S3. Cellular GWAS of mean IAV burden in LCLs.** (A) Manhattan plot of mean IAV burden based on p values from EMMAX using kinship matrix, sex as a covariate, and with SNPs with MAF < 10% and deviation from Hardy-Weinberg Equilibrium (p < 1x10-4) excluded. P values plotted are corrected for genomic inflation factor ($\lambda = 1.05$). (B) QQ plot of the same data in A.

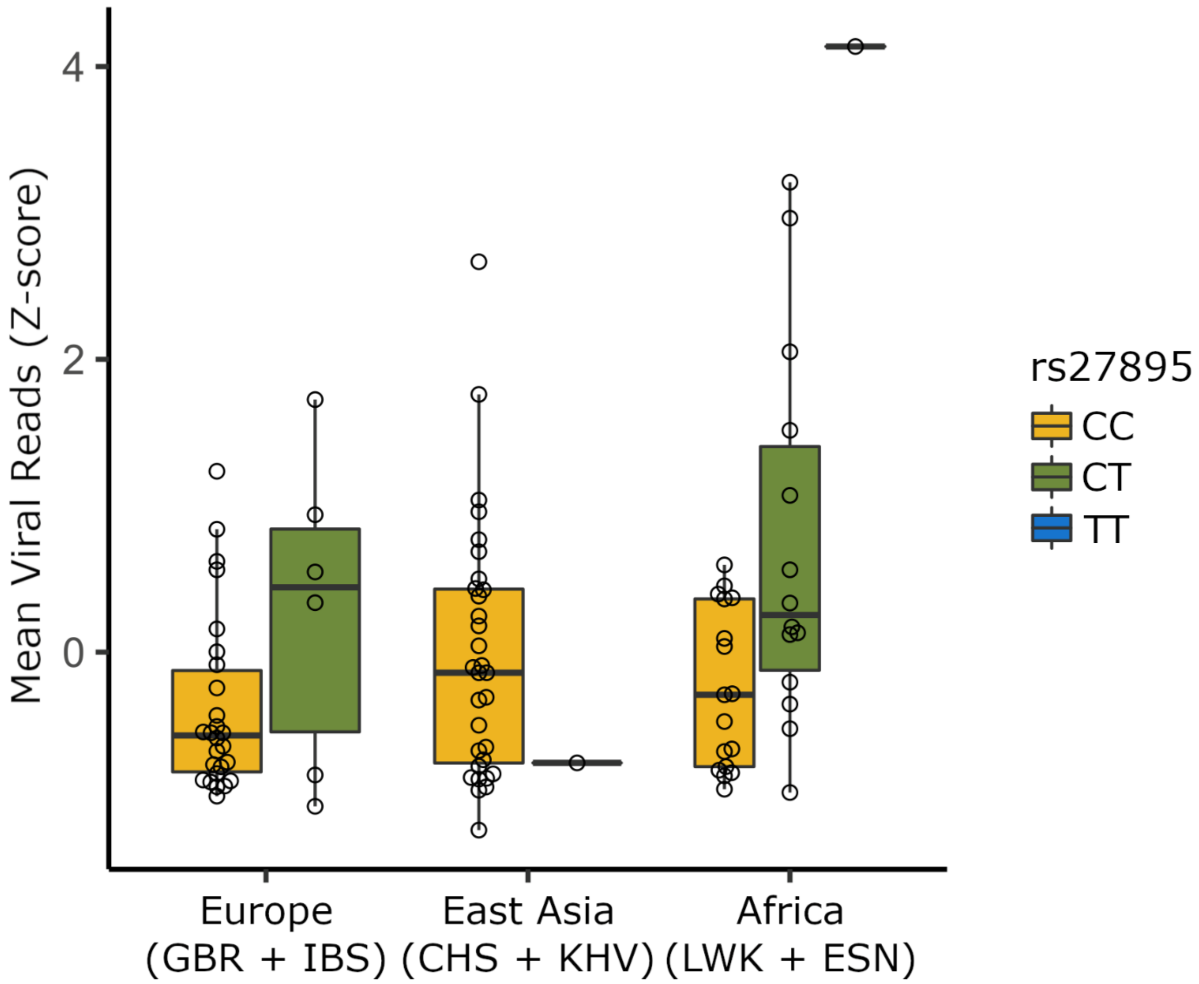


**Figure S4. Genotype median plot for rs27895 mean IAV reads in individual scHi-HOST populations.** Plotting by continent reveals that rs27895-T is associated with increased viral reads in European and African populations while the C allele of rs27895 is nearly fixed in East Asian populations.

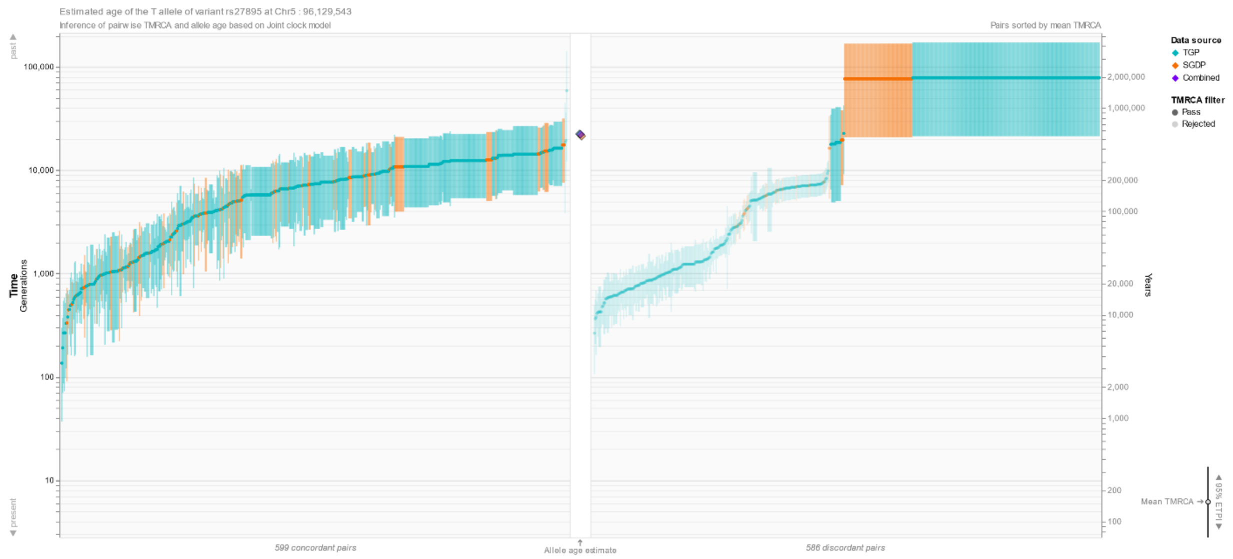


**Figure S5. Estimate of the age of rs27895 suggests that the variant was present in human populations before the out-of-Africa expansion.** Data and figure from https://human.genome.dating
